# The glycolipid GlcCer is recruited into the viral envelope to promote phenuivirus binding to host cells

**DOI:** 10.1101/2022.08.07.502911

**Authors:** Zina M. Uckeley, Magalie Mazelier, Christian Lüchtenborg, Sophie L. Winter, Paulina Schad, Petr Chlanda, Britta Brügger, Pierre-Yves Lozach

## Abstract

Virus–receptor interactions largely contribute to the tropism and outcome of an infection. Here, we found that the glycolipid glucosylceramide (GlcCer) is a major component of Uukuniemi phenuivirus and allows viral binding to host cells. A lipidomic analysis with mass spectrometry revealed the lipidome of UUKV particles and indicated that GlcCer was enriched in both infected cells and viral particles. In addition, the infectivity of UUKV depended on the conversion of ceramide (Cer) into GlcCer in the Golgi network of producer cells. In contrast, depletion of GlcCer in virions profoundly impaired the attachment of UUKV and other related viruses to target cells. Furthermore, competing GlcCer ligands prevented virus binding to various cell types. Altogether, our results demonstrate that glycolipids are essential structural determinants of the virions necessary for virus attachment to host cells and have strong implications for future work on the identification of virus receptors.

## Introduction

*Phenuiviridae* in the *Bunyavirales* order is a large family of RNA viruses that comprises 19 genera and more than 100 members worldwide (Koch et al., 2021). These viruses represent an emerging global threat to human public health and agricultural productivity (Peyre et al., 2015). Many phenuivirus species are causes of birth defects and serious diseases in both humans and livestock, such as acute hepatitis, hemorrhagic fever, and encephalitis. Uukuniemi virus (UUKV) is used as a viral model system to study the closely related Dabie virus (DABV), previously referred to as severe fever with thrombocytopenia syndrome virus (SFTSV), and other highly pathogenic phenuiviruses that have recently emerged in China and North America (Rezelj et al., 2015). Infection with another phenuivirus, Toscana virus (TOSV), is the primary cause of arboviral diseases in humans in southern Europe during the summer season (Hemmersbach-Miller et al., 2004; Terrosi et al., 2009; Vilibic-Cavlek et al., 2020). Phenuiviruses are spread mainly by hematophagous arthropods and, hence, belong to the supergroup of arthropod-borne viruses (arboviruses). Owing to their mode of transmission, global warming, and the increasing number of recent outbreaks worldwide, phenuiviruses are considered potential agents of emerging diseases (Elliott and Brennan, 2014). To date, no vaccines or specific antiviral treatments have been approved for human use.

At the molecular level, phenuiviral particles are enveloped and roughly spherical (approximately 100 nm in diameter) with a trisegmented single-stranded RNA genome that replicates exclusively in the cytosol of infected cells. The viral structural proteins are encoded in the negative sense orientation with the nucleoprotein N encoded in the small genomic segment (S), the two transmembrane envelope glycoproteins Gn and Gc encoded in the medium segment (M), and the RNA-dependent RNA polymerase (RdRp) encoded in the large segment (L) (Koch et al., 2021). The viral particles assemble and bud in the Golgi, where they acquire a lipid bilayer envelope and exit from infected cells (Overby et al., 2006).

The surface of a virion in the phenuivirus family is decorated by the glycoproteins Gn and Gc, which are arranged in a T=12 icosahedral symmetry and are critical for viral penetration into the cytosol (Freiberg et al., 2008; Överby, 2008). Structural studies indicated that Gc is a class II fusion protein. Within virions, the N protein is associated with the RNA genome and, together with the viral RdRp, forms three ribonucleoproteins (RNPs).

The transmission, tropism, and cellular receptors leveraged by phenuiviruses to enter and infect host cells have not been extensively characterized. Heparan sulfates have been shown to facilitate TOSV and Rift Valley fever virus (RVFV) infections (Boer et al., 2012; Pietrantoni et al., 2015; Riblett et al., 2016), and nonmuscle myosin heavy chain type-IIA (NMMHC-IIA) has been proposed to be an attachment factor for DABV (Sun et al., 2014). The human C-type lectins dendritic cell-specific intercellular adhesion molecule 3-grabbing non-integrin (DC-SIGN) and liver/lymph node cell-specific intercellular adhesion molecule 3-grabbing non-integrin (L-SIGN) have been identified as entry receptors for RVFV, TOSV, Punta Toro virus (PTV), and UUKV (Lozach et al., 2011; Léger et al., 2016). These lectins have been suggested to play roles in the tropism of these viruses in dermal dendritic cells (DCs) and liver cells, respectively (Koch et al., 2021). DC-SIGN is located on the surface of dermal DCs, while L-SIGN is present on the endothelial lining of hepatic sinusoids. It is however apparent that phenuiviruses likely rely on other receptors to target and bind host cells since they have a broad tissue tropism, most of which do not express DC-SIGN or L-SIGN. Upon attachment, phenuiviruses are sorted into the endocytic machinery and penetrate host cells via acid-activated membrane fusion (Koch et al., 2021).

Interestingly, viral envelope lipids have been reported to modulate binding and thus regulate the tropism of unrelated viruses to a host. Transmembrane immunoglobulin and mucin domain (TIM) and Tyro3, Axl, and Mer (TAM) proteins, two families of receptors mediating phosphatidylserine (PS)-dependent phagocytosis, are receptors of flaviviruses and filoviruses (Amara and Mercer, 2015). Although a specific interaction between PS in viral envelopes and TIM/TAM remains to be identified, previous observations strongly suggest that viral envelope lipids are important binding modulators. Overall, the lipid composition of enveloped viruses and their functional significance remain to be sufficiently characterized, and little information is available on the importance of lipids to phenuivirus entry.

To examine the roles played by lipids in early phenuivirus–host cell interactions, we subjected UUKV particles produced in mammalian tissue culture cells to a lipidomic analysis with mass spectrometry (MS). Using this approach, we identified the glycolipid glucosylceramide (GlcCer) as a crucial viral factor for the attachment of UUKV and other bunyaviruses to various cell types. This work opens the way to the study of a newly discovered type of virus–receptor interaction: the interaction between glycolipids in the viral envelope and receptors on the host cell surface.

## Results

### A quantitative lipidomic analysis led to the identification of hexosylceramide (HexCer) as a major component of the UUKV particles

The lipid composition of phenuiviruses have not been extensively documented. To better understand the role played by lipids in the viral envelope on phenuivirus–host cell interactions, we first employed a lipidomic analysis with MS approach to determine the lipid distribution in cells after UUKV infection. To this end, Baby hamster kidney fibroblasts (BHK-21 cells) were exposed to UUKV and harvested 48 h post-infection, after which several cycles of infection have occurred and virus progeny production is peaking (Lozach et al., 2010). BHK-21 cells constitute the gold standard model system to study UUKV and other phenuiviruses, and in these cells, a complete UUKV life cycle, from infection to release of infectious progeny, is completed in approximately 7 h (Lozach et al., 2010).

Infected cells were subjected to a quantitative MS-based lipid analysis applying a shotgun lipidomic approach (Vvedenskaya et al., 2021). We quantitatively assessed 374 lipid species in 22 lipid classes, including the glycerophospholipids phosphatidylcholine (PC), lyso-PC (LPC), phosphatidylethanolamine (PE), phosphatidylinositol (PI), PS, phosphatidic acid (PA), PE plasmalogen (PE P), and phosphatidylglycerol/lysobisphosphatidic acid (PG/LBPA) (Table S1). PC, PE, PS, PI, and PG were further subcategorized into diacyl designated without a prefix, indicating the type and species with either plasmanyl/acyl or diacyls, and with the prefix O, indicating even/odd chain fatty acyl composition. A sphingolipid species analysis was performed to detect sphingomyelin (SM), ceramide (Cer), HexCer, and dihexosylceramide (Hex2Cer) species. The sterol lipids identified included cholesterol (Chol) and cholesteryl ester (CE) species, and triacylglycerol (TAG) and diacylglycerol (DAG) were representative neutral lipids. The results showed that infection did not affect the distribution of the lipid classes in producer cells, with the exception of HexCer, the level of which was increased 5-fold compared to that of uninfected cells (Figure 1A, Table S1). HexCers comprises GlcCer and galactosylceramide (GalCer) that are glycolipids in the endoplasmic reticulum (ER) and Golgi apparatus, where they are transient intermediates in glycosphingolipid (GSL) synthesis (Figure 1B) (Burger et al., 1996). The MS analysis revealed that the abundance of Cer and Hex2Cer, upstream and downstream intermediates of HexCer in this GSL producing pathway, was not affected by viral infection, as indicated by the levels of both lipids remaining at a minimal level (Figure 1A). Taken together, our results showed that UUKV infection leads to HexCer accumulation in cells, presumably in the Golgi apparatus.

**Figure 1.**
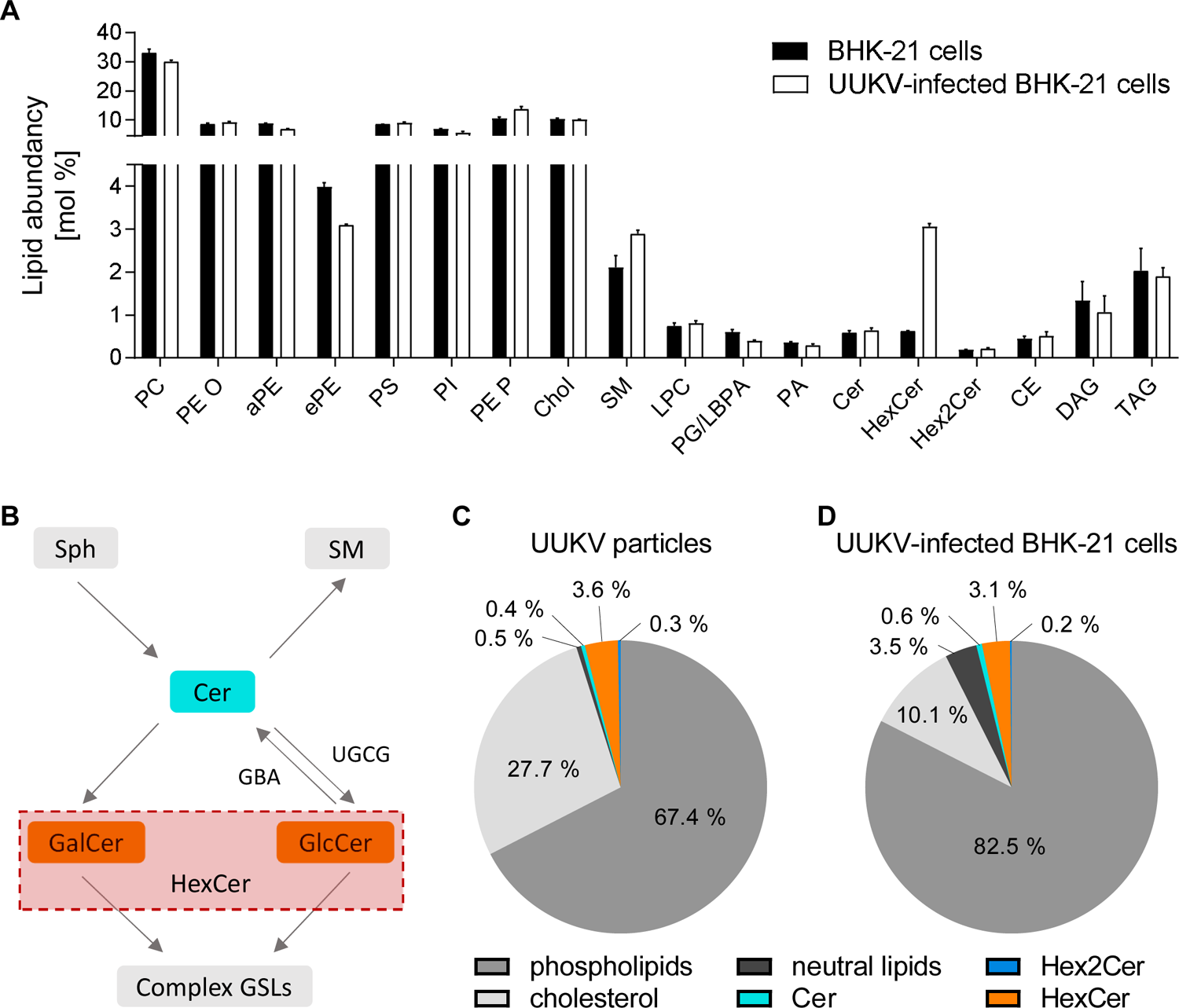
A label-free quantitative lipidomic analysis led to the identification of hexosylceramide (HexCer) as a major component of Uukuniemi virus (UUKV) particles. (A) BHK-21 cells were infected with UUKV at a multiplicity of infection of 0.1 for 48 h and then subjected to a lipidomic analysis. Lipid classes are presented as mol % fractions. (B) Schematic overview showing HexCer metabolism. Cer, ceramide; GBA, β-glucocerebrosidase; GalCer, galactosylceramide; GlcCer, glucosylceramide; GSL, glycosphingolipid; SM, sphingomyelin; Sph, sphingosine; UGCG, UDP-GlcCer glucosyltransferase. (C) Supernatant from infected BHK-21 cells was harvested 48 h post-infection, and UUKV particles were purified prior to lipid MS analysis. (D) Lipidome of UUKV-infected BHK-21 cells. (C and D) The phospholipids included all glycerophospholipids such as phosphatidylethanolamine plasmalogen (PE P), SM, and lysophosphatidylcholine (LPC). The neutral lipids were cholesteryl ester (CE), diacylglycerol (DAG), and triacylglycerol (TAG).

The Golgi network is the assembly site of UUKV and other phenuiviruses (Overby et al., 2006). We, therefore, wondered whether an increase in HexCer levels leads to its incorporation into the envelope of UUKV particles. To assess this possibility, we performed a lipidomic analysis with MS of purified viral particles produced in BHK-21 cells (Figure 1C, Figure S1A, and Table S2). In these experiments, we additionally used virus-free mock cell supernatants to control for lipid contaminants of the UUKV preparations. No significant amount of lipid was present in the virus-free mock cell supernatant after the purification process. We found that Chol and the phospholipids SM and PE P were enriched in UUKV particles compared to the total lipid composition in the producer cells (Figure 1A, 1C, 1D, and S1A and Table S1 and S2). These lipid classes are typical of the post-*cis* stacks of the Golgi network (Surma et al., 2011; Duran et al., 2012), consistent with budding of viral particles from this compartment. HexCers accounted for almost 4% of the lipids in the viral envelope (Figure 1C).

Overall, our data indicated that UUKV infection modulates the GSL synthesis pathway such that HexCer is both enriched in infected cells and is incorporated into viral particles during budding. In all further experiments, we sought to determine whether the incorporation of HexCer into the viral envelope exhibited a specific function in phenuivirus–host cell interactions.

### UUKV depends on GlcCer synthase and GlcCer for infection and spread

Our lipidomic analysis with MS approach did not allow us to discriminate the nature of the HexCer molecules incorporated into the UUKV particles, *i.e.*, GlcCer or GalCer (Figure 1 and S1A and Table S1 and S2). As GlcCer is the most abundant HexCer in a broad spectrum of tissues, we focused on GlcCer and assessed its potential involvement in the spread of UUKV in tissue culture. We first aimed to disrupt Cer conversion into GlcCer (Figure 1B) by reducing the expression of UDP-GlcCer glucosyltransferase (UGCG) with two in-house-designed, nonoverlapping small interfering RNAs (siRNAs) in parallel experiments. With the application of each siRNA, the level of UGCG in the cultures was reduced by 60-80%, as determined by western blotting using an antibody targeting UGCG (Figure 2A and 2B). The silencing of UGCG expression resulted in a decrease in GlcCer synthesis by approximately 60-80%, as shown by dot blotting with an antibody against GlcCer (Figure 2C and 2D). We then exposed cells with silenced UGCG to UUKV at a multiplicity of infection (MOI) of 0.1 for 24 h. The sensitivity of these cells to UUKV infection was assessed by flow cytometry analysis after immunostaining for the viral nucleoprotein N (Figure 2E). Approximately 35-40% of cells transfected with scrambled siRNA were found to be UUKV N-positive in this assay. When UGCG was silenced, UUKV infection was reduced by 60-70% in cells carrying either UGCG siRNA compared to that in the control cells transfected with nontargeted control siRNA (Figure 2E and 2F).

**Figure 2.**
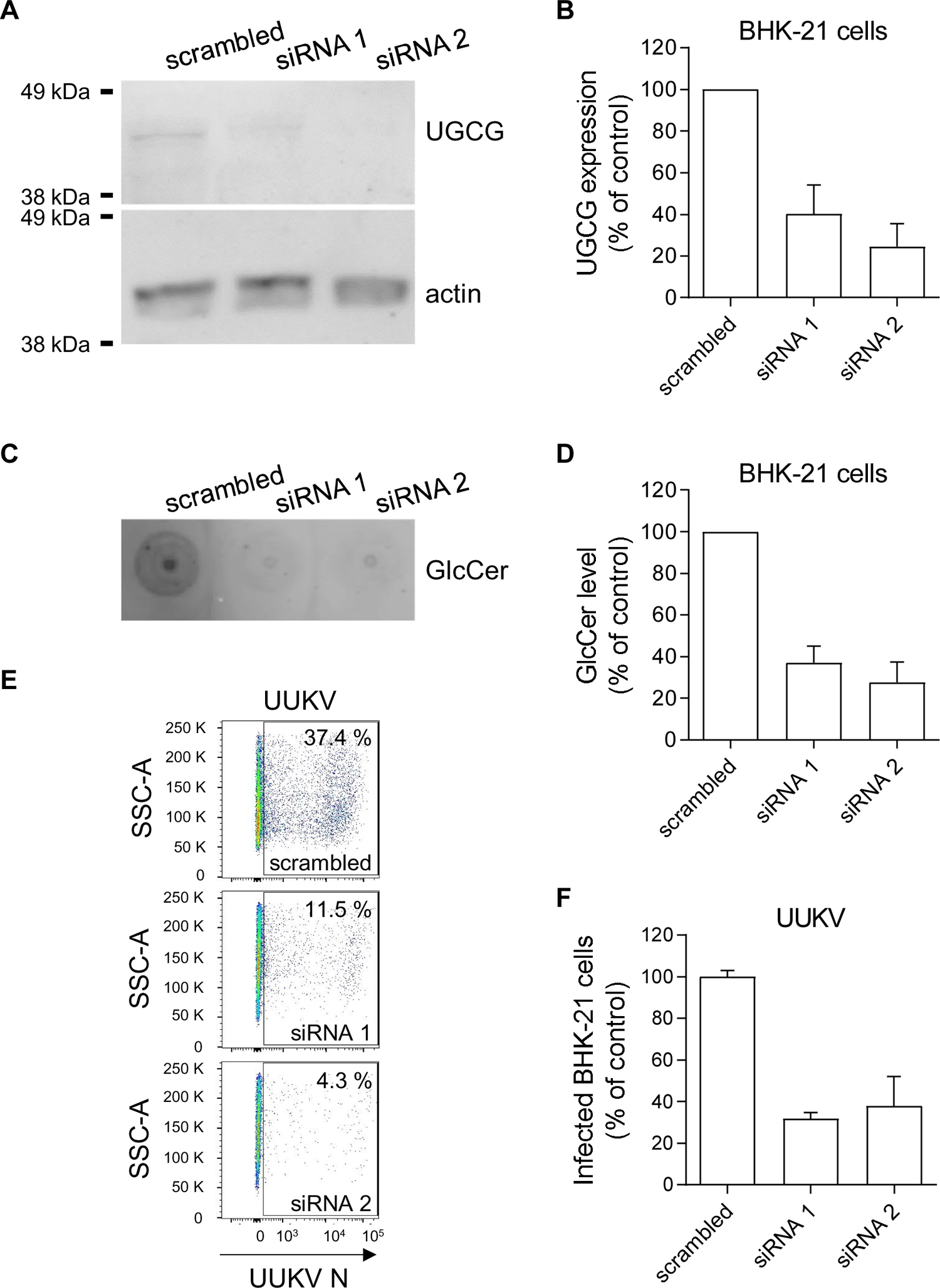
Glucosylceramide (GlcCer) synthase (UGCG) depletion inhibits Uukuniemi virus (UUKV) infection. (A) UGCG synthase was silenced in BHK-21 cells with two non-overlapping short interfering RNAs (siRNAs; 20 nM) and assayed by western blotting 72 h later. (B) The efficiency of UGCG knockdown was semiquantified under the conditions described in (A). UGCG protein levels are reported as the percentage of UGCG levels in cells treated with siRNAs against UGCG and normalized to levels of actin and UGCG in BHK-21 cells treated with negative-control siRNAs (scrambled). (C) GlcCer levels were examined in BHK-21 cells after silencing GlcCer synthase with siRNAs by dot blotting. (D) GlcCer levels were semiquantified on the basis of the dot blots shown in (C) and expressed as a percentage of GlcCer levels in cells treated with siRNAs against GlcCer and normalized to GlcCer levels in BHK-21 cells treated with negative control siRNAs (scrambled). (E) BHK-21 cells were treated with siRNAs against UGCG (20 nM) for 72 h and then exposed to UUKV (multiplicity of infection ∼0.1). Infection was determined by flow cytometry after immunostaining for the viral nucleoprotein N 24 h post-infection. (F) The values were normalized to the infection level in samples treated with siRNA controls (scrambled).

We next assessed the ability of chemical inhibitors of UGCG to prevent UUKV infection. We first treated cells with DL-*threo*-1-phenyl-2-palmitoylamino-3-morpholino-1-propanol (PPMP), a Cer analog (Abe et al., 1992; Atilla-Gokcumen et al., 2011; Alam et al., 2015). We found that the maximal concentration for which PPMP exerted no adverse effects on BHK-21 cell viability was 5 µM, as determined with a quantitative assay measuring the release of lactate dehydrogenase (LDH) into the extracellular medium upon cell death and lysis (Figure S2). The distribution of the major classes of select lipids, including Cer and Hex2Cer, was not affected in the UUKV-infected BHK-21 cells that were treated with PPMP at concentrations as high as 5 µM for 24 h (Figure 3A and 3B and Table S3). Only the PG/LBPA was enriched, consistent with the accumulation of LBPA in endosomal/lysosomal structures, as has recently been reported for such treatment (Hartwig and Höglinger, 2021). Under these conditions, PPMP abolished the increase in HexCer in infected cells (Figure 3B). A dot blot analysis confirmed that the GlcCer level was decreased by 70-80% in cells after treatment with PPMP (Figure 3C and 3D). Taken together, these data demonstrated that PPMP hampered UGCG and, in turn, GlcCer synthesis and HexCer enrichment in UUKV-infected cells.

**Figure 3.**
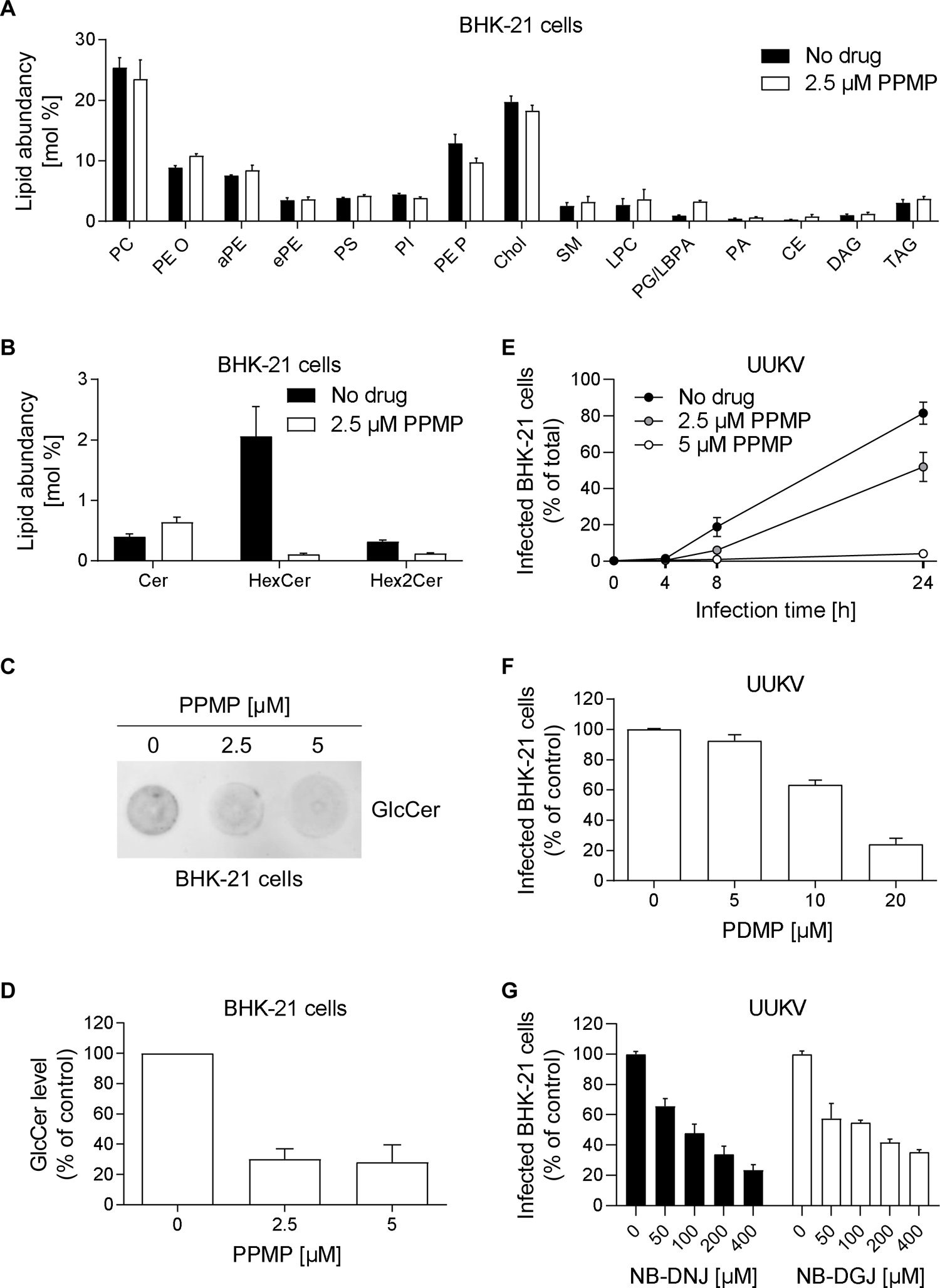
Uukuniemi virus (UUKV) relies on glucosylceramide (GlcCer) in viral particles for cell entry. (A) BHK-21 cells were infected with UUKV (multiplicity of infection (MOI) ∼0.1) in the presence of DL-*threo*-1-phenyl-2-palmitoylamino-3-morpholino-1-propanol (PPMP), which inhibits UGCG activity. Infected cells were subjected to lipidomics analyses 24 h post-infection. (B) Specific analysis of HexCer and Hex2Cer levels in the samples processed as described in (A). (C) GlcCer levels were analyzed in BHK-21 cells after PPMP treatment by dot blotting. (D) GlcCer levels in (C) were semiquantified and expressed as a percentage of the GlcCer level measured in the absence of PPMP. (E) BHK-21 cells were first treated with PPMP for 16 h and then infected with UUKV (MOI ∼0.5) in the continuous presence of the drug. Infection was detected by flow cytometry up to 24 h later. Note that the SEMs of some data series are not visible on the graph. (F and G) BHK-21 cells were pretreated with three other UGCG inhibitors, namely, N-[2-hydroxy-1-(4-morpholinylmethyl)-2-phenylethyl]-decanamide (PDMP), N-Butyldeoxynojirimycin (NB-DNJ), or N-Butyldeoxygalactonojirimycin (NB-DGJ) at the indicated concentrations for 16 h (PDMP) or 24 h (NB-DNJ and NB-DGJ) and were then infected with UUKV (MOI ∼0.1) in the continuous presence of the drugs. Infection was quantified by flow cytometry after immunostaining for the viral nucleoprotein, and the data were normalized to those in control samples without inhibitor treatment.

To assess the sensitivity of UUKV infection to PPMP, BHK-21 cells were infected at an MOI of 0.1 in the continuous presence of PPMP for up to 24 h. Using our flow cytometry-based infection assay, we observed that PPMP reduced UUKV infection in a dose-dependent manner (Figure 3E). To exclude the possibility that this result was cell line-specific, we assessed the effect of PPMP treatment on UUKV infection in the A549 human lung epithelial cell line, which is susceptible for UUKV infection (Lozach et al., 2010). In the A549 cells, PPMP administered in concentrations ranges identical to those administered to BHK-21 cells did not induce cytotoxicity (Figure S2). Similar to the results obtained with BHK-21 cells, pretreatment of the A549 cells with PPMP blocked UUKV infection (Figure S3).

Through complementary approaches, we examined three other UGCG inhibitors, namely, N-[2-hydroxy-1-(4-morpholinylmethyl)-2-phenylethyl]-decanamide (PDMP), N-butyl-deoxynojirimycin (NB-DNJ), and N-butyl-deoxygalactonojirimycin (NB-DGJ), for their capacity to prevent UUKV infection. PDMP is another Cer analog that is similar to PPMP, while NB-DNJ and NB-DGJ are both glycan analogs that compete with glucose to inhibit UGCG activity (Andersson et al., 2000; Grabowski, 2008). BHK-21 cells were treated with each of these drugs and then infected with UUKV, and the results were similar to those obtained with PPMP. The three drugs each negatively affected UUKV infection in a dose-dependent manner (Figure 3F and 3G). Collectively, our results showed that UUKV relies on UGCG and GlcCer for infection and likely spread. All inhibitors displayed similar efficiency in preventing UUKV infection, and we used PPMP in the following experiments.

### UUKV progeny relies on GlcCer for infectivity

Next, we sought to determine the steps in the UUKV life cycle in cells that were impaired when UGCG activity was inhibited or GlcCer was depleted. To this end, we harvested the supernatant of UUKV-infected BHK-21 cells treated with 2.5 µM PPMP. Then, the supernatant was used to infect freshly seeded BHK-21 cells for 8 h to ensure a single round of infection. We observed that the infection of the cells cultured in supernatant from PPMP-treated infected cells was reduced by more than 70% when compared to the infection of cells exposed to the supernatant from mock-treated infected cells (Figure 4A). We then measured infectious UUKV progeny in the supernatant of GlcCer-depleted producer BHK-21 cells. Performing a standard foci-forming unit titration assay (Lozach et al., 2010), we found that the number of infectious UUKV particles dropped by 90% in the supernatant of the infected cells exposed to PPMP compared with the untreated cells (Figure 4B). The amount of UUKV proteins, which reflects the number of viral particles, was reduced by 50% in the presence of PPMP, as determined by western blotting (Figure 4C and 4D). This indicated that virion production is partially inhibited in the presence of PPMP. The decrease in the total number of viral particles, however, did not seem to completely reflect the decrease in infectivity of GlcCer-depleted virions (Figure 4B and 4D).

**Figure 4.**
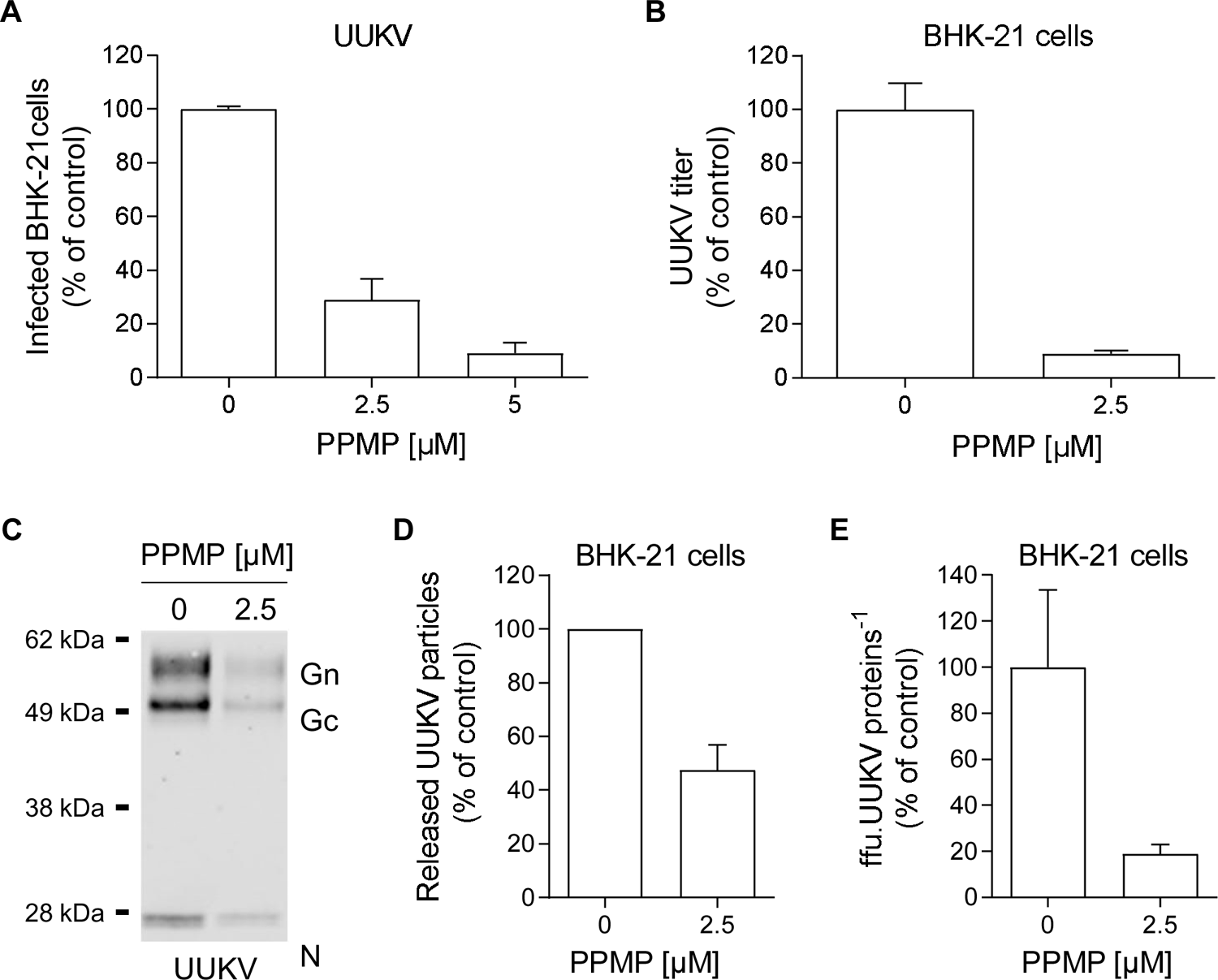
Glucosylceramide (GlcCer) is critical for Uukuniemi virus (UUKV) infectivity. (A) BHK-21 cells were pretreated with DL-*threo*-phenyl-2-palmitoylamino-3-morpholino-1-propanol (PPMP) at the indicated concentrations for 16 h and exposed to UUKV (multiplicity of infection ∼0.1). Supernatants of infected cultured cells were harvested 24 h post-infection and then allowed to infect freshly seeded naïve BHK-21 cells for 8 h. The infected cells were immunostained for detection of UUKV N and analyzed by flow cytometry. The values were normalized to those in control samples where viruses were produced in the absence of the inhibitor. (B) Infectious UUKV progeny in the supernatant collected from cells cultured in the presence of PPMP was evaluated by focus-forming assay. The data are presented as the percentage of the control samples where viruses were produced in the absence of the inhibitor. (C) Viral particles produced from BHK-21 cells treated with 2.5 µM of PPMP were analyzed by SDS–PAGE and western blotting under nonreducing conditions with a polyclonal antibody that recognized UUKV N, Gn, and Gc. (D) UUKV structural proteins in (C) were semiquantified. The data were expressed as a percentage of the level of UUKV N, Gn, and Gc in the supernatant of PPMP-treated and infected BHK-21 cells and normalized to the level of N, Gn, and Gc in the supernatant of infected cells that had not been exposed to PPMP. (E) The ratio of the number of focus-forming units (ffus) to the relative unit of UUKV structural proteins is presented as the percentage of the value obtained for the control sample without any drug treatment.

To clarify whether PPMP exerted an impact on the infectivity of individual viral particles or on the number of total UUKV particles produced in cells, we determined the ratio of infectious particles to total particles, *i.e.*, the ratio of infectious particles to the total amount of UUKV proteins. This ratio was reduced by 80% in the medium of PPMP-treated cells, indicating that the infectivity of UUKV progeny was profoundly compromised in the absence of GlcCer (Figure 4E). Overall, the results highlighted the importance of GlcCer for the infectivity of viral particles released into the medium and the early steps of UUKV infection.

### The absence of GlcCer does not impact the structure and protein composition of UUKV particles

Host cell receptors have been shown to bind phenuivirus Gn and Gc envelope glycoproteins (Koch et al., 2021). We therefore evaluated the impact of GlcCer depletion induced by PPMP treatment on the amount of Gn and Gc embedded in viral particles. For this purpose, we determined the ratio of UUKV glycoproteins to N proteins in purified viral particles as analyzed by western blotting (Figure 4C). This showed no significant difference in the UUKV glycoprotein-to-N ratio whether GlcCer was present in the viral envelope or not (Figure 5A).

**Figure 5.**
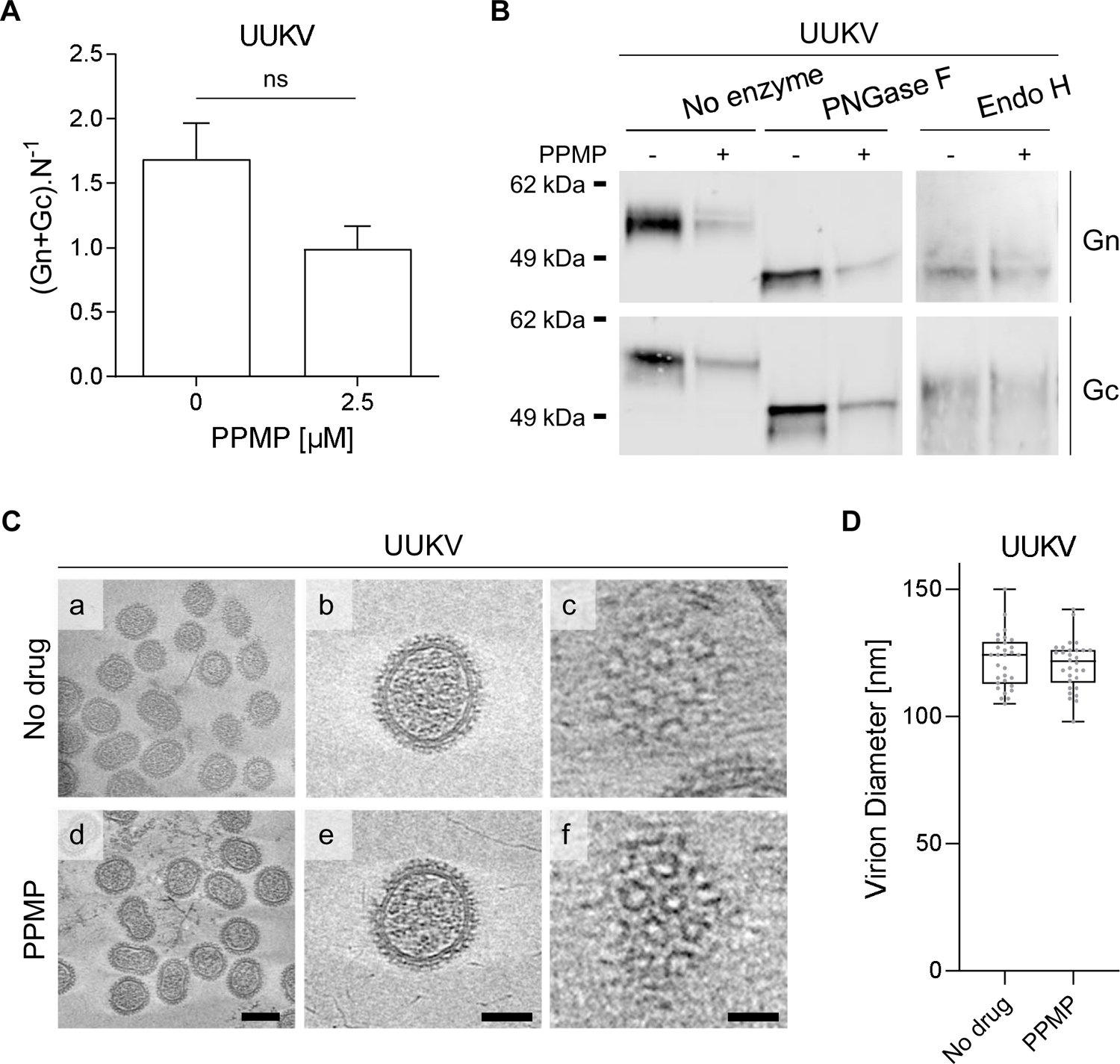
Uukuniemi viruses (UUKVs) with and without glucosylceramide (GlcCer) show no morphological differences. (A) UUKV particles produced from BHK-21 cells in the presence of DL-*threo*-phenyl-2-palmitoylamino-3-morpholino-1-propanol (PPMP, 2.5 µM) were analyzed and semiquantified by western blotting as described in Figure 4C and 4D. Values are shown as the ratio of the UUKV glycoproteins Gn and Gc level to nucleoprotein N level. (B) UUKV derived from BHK-21 cells exposed to PPMP was digested with PNGase F (1,000 units) or Endo H (2,000 units) under reducing conditions for 4 h at 37°C. Proteins were separated by SDS–PAGE and analyzed by western blotting with a mAb against either UUKV glycoproteins Gn or Gc. ns, not significant. (C) Cryo-electron tomography of gradient purified UUKV in the presence or absence of PPMP. (a, d) Slices of tomogram capturing UUKV virions. Scale bars correspond to 100 nm (a, d) and 50 nm (b, e). (c, f) Tomograph slices showing Gn and Gc virion surface arrangement (scale bar: 20 nm). (D) The diameter of UUKV particles produced in the presence or absence of PPMP was measured (n=30).

The UUKV glycoproteins Gn and Gc carry several *N*-linked oligosaccharides (four glycosylation sites in each) that contribute to the direct binding of UUKV to C-type lectin receptors (Lozach et al., 2011; Léger et al., 2016). Hence, we analyzed the glycosylation pattern of UUKV Gn and Gc in the viral particles produced in cells depleted from GlcCer because of PPMP treatment. UUKV particles were treated with *N*-glycosidase F (PNGase F) and endoglycosidase H (Endo H) before separation by SDS–PAGE and western blotting (Figure 5B). Both viral glycoproteins were found to be sensitive to PNGase F and Endo H to a similar extent, whether or not GlcCer was embedded in the viral envelope. The data indicated that *N*-glycan residues in UUKV Gn and Gc do not account for the defective binding of GlcCer-depleted UUKV particles.

Finally, to confirm that PPMP does not impact the overall structural organization of UUKV particles, we imaged viral preparations via cryo-electron tomography (cryo-ET). No apparent difference was observed between the virions produced in the absence or presence of PPMP, indicating that PPMP exerted no impact on virus assembly (Figure 5C, panels a to f). The typical arrangement of Gn and Gc was visible on the surface of the viral particles regardless of the cell treatment (Figure 5C, panels c and f), and no difference in virion size was detected (Figure 5D). Overall, these results suggest that depletion of GlcCer from the viral envelope does not alter the assembly and structure of UUKV.

### GlcCer in the viral envelope promotes UUKV binding to target cells

Our results suggested that UUKV relies on GlcCer for infectious entry. To confirm this possibility, we evaluated the role played by GlcCer in virus binding to BHK-21 cells. To this end, the amount of input virus was normalized based on the UUKV N protein level in the virus stocks as measured by western blotting and not the MOI. The N level reflects the total number of viral particles, in contrast to the MOI, which reveals only infectious virus levels. UUKV particles were produced in the presence or absence of PPMP for 24 h. Fresh, naïve BHK-21 cells were then exposed to virions on ice for 2 h, extensively washed, and subjected to SDS– PAGE and western blot analysis (Figure 6A). Strikingly, the amount of UUKV nucleoprotein N bound to the BHK-21 cells was reduced by 60-70% when the viral particle envelopes lacked GlcCer (Figure 6A and 6B).

**Figure 6.**
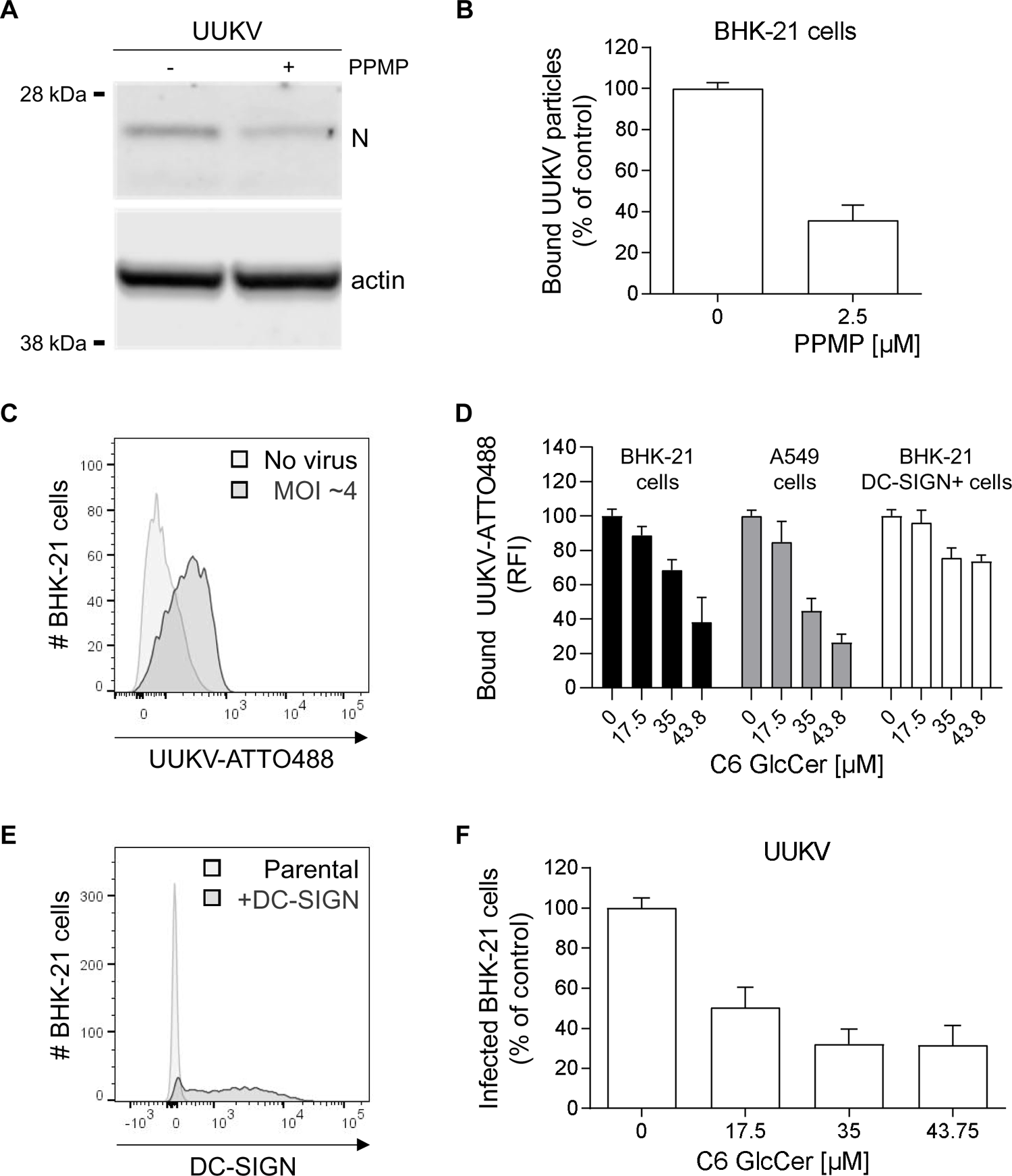
Glucosylceramide (GlcCer) in viral particles promotes Uukuniemi virus (UUKV) binding. (A) UUKV particles derived from BHK-21 cells in the presence of DL-*threo*-phenyl-2-palmitoylamino-3-morpholino-1-propanol (PPMP, 2.5 µM) were bound to freshly seeded naïve BHK-21 cells for 2 h on ice before fixation and western blot analysis with an antibody recognizing the UUKV N protein. (B) N was semiquantified from the cells described in (A), and the value is presented as a percentage of the N level measured in the sample corresponding to virus binding in the absence of PPMP. (C) Fluorescently labeled UUKV particles (UUKV-ATTO488) were bound to BHK-21 cells (multiplicity of infection (MOI) ∼4) on ice for 1 h, and viral binding was evaluated by flow cytometry analysis. (D) BHK-21 cells, A549 human lung epithelial cells, and BHK-21 cells expressing the UUKV receptor DC-SIGN (BHK-21 DC-SIGN+) were preincubated with varying amounts of soluble C6-GlcCer for 2 h and then exposed to UUKV-ATTO488 (MOI ∼4) on ice for 1 h. Virus binding was measured by flow cytometry, and the data were normalized to those in control samples processed in the absence of soluble C6-GlcCer. RFI, relative fluorescence intensity. (E) BHK-21 cells were transduced with a retroviral vector system to express DC-SIGN (BHK-21 DC-SIGN+). DC-SIGN expression was measured by flow cytometry analysis using phycoerythrin-conjugated anti-DC-SIGN mAb. (F) Soluble C6-GlcCer was allowed to bind BHK-21 cells on ice for 2 h prior to exposure to UUKV on ice for 1 h (MOI ∼0.5). After virus binding on ice, unbound UUKV particles were washed away, and the cells were incubated at 37°C for 8 h. Infection was quantified by flow cytometry after immunostaining for UUKV N protein. Values are presented as the percentage of the control sample without prebinding of soluble C6-GlcCer.

To further investigate the possibility that GlcCer directly promotes the binding of viral particles to host cells, we subjected ATTO488-conjugated UUKV (UUKV-ATTO488) to a flow cytometry-based binding competition assay (Figure 6C) (Lozach et al., 2011; Hoffmann et al., 2018). Binding of fluorescently labeled UUKV particles to BHK-21 and A549 cells was found to be inhibited by prebinding of soluble C6-GlcCer, and the effect was dose-dependent (Figure 6D). In contrast, DC-SIGN expression in BHK-21 cells, which do not endogenously express this lectin (Lozach et al., 2011), largely preserved UUKV binding in the presence of prebound C6-GlcCer (Figures 6D and 6E). This finding indicated that UUKV binding can occur through interactions between DC-SIGN and mannose residues in the Gn and Gc glycoproteins or GlcCer present in the viral envelope and an unknown receptor in BHK-21 cells. In addition, binding competition between soluble C6-GlcCer and UUKV before incubation at 37°C significantly reduced viral infection (Figure 6F). Taken together, these experiments indicated that UUKV binding to BHK-21 and A549 cells is mediated through GlcCer in the viral envelope and that binding most likely involves specific attachment factors or receptors but not DC-SIGN.

### GlcCer is necessary for infectious entry of phenuiviruses and other bunyaviruses

Next, we examined whether other viruses depend on GlcCer to bind to target cells. First, we analyzed BHK-21 cells via our lipidomic with MS approach after the cells were infected with Semliki forest virus (SFV), an unrelated virus from the *Togaviridae* family that assembles at the plasma membrane (Kai Simons, 1980; Simons and Warren, 1984). SFV did not alter the distribution of Cer, HexCer, or Hex2Cer in infected cells (Figure 7A and Table S4), suggesting that infection by SFV does not interfere with the GSL synthesis pathway. Next, SFV particles produced by these cells were subjected to lipid MS analysis after purification, and the results also did not indicate increased GlcCer incorporation into the virions (Figure 7B and S1B, and Table S2). Identical results were obtained with Ebola virus-like particles (EBOVLPs), which also buds from the plasma membrane (Figure 7C). The total amount of Cer and all its derivatives did not exceed 2.6% of the EBOVLP envelope; in contrast, 3.6% of the UUKV envelope consists of HexCer alone (Figure 1C).

**Figure 7.**
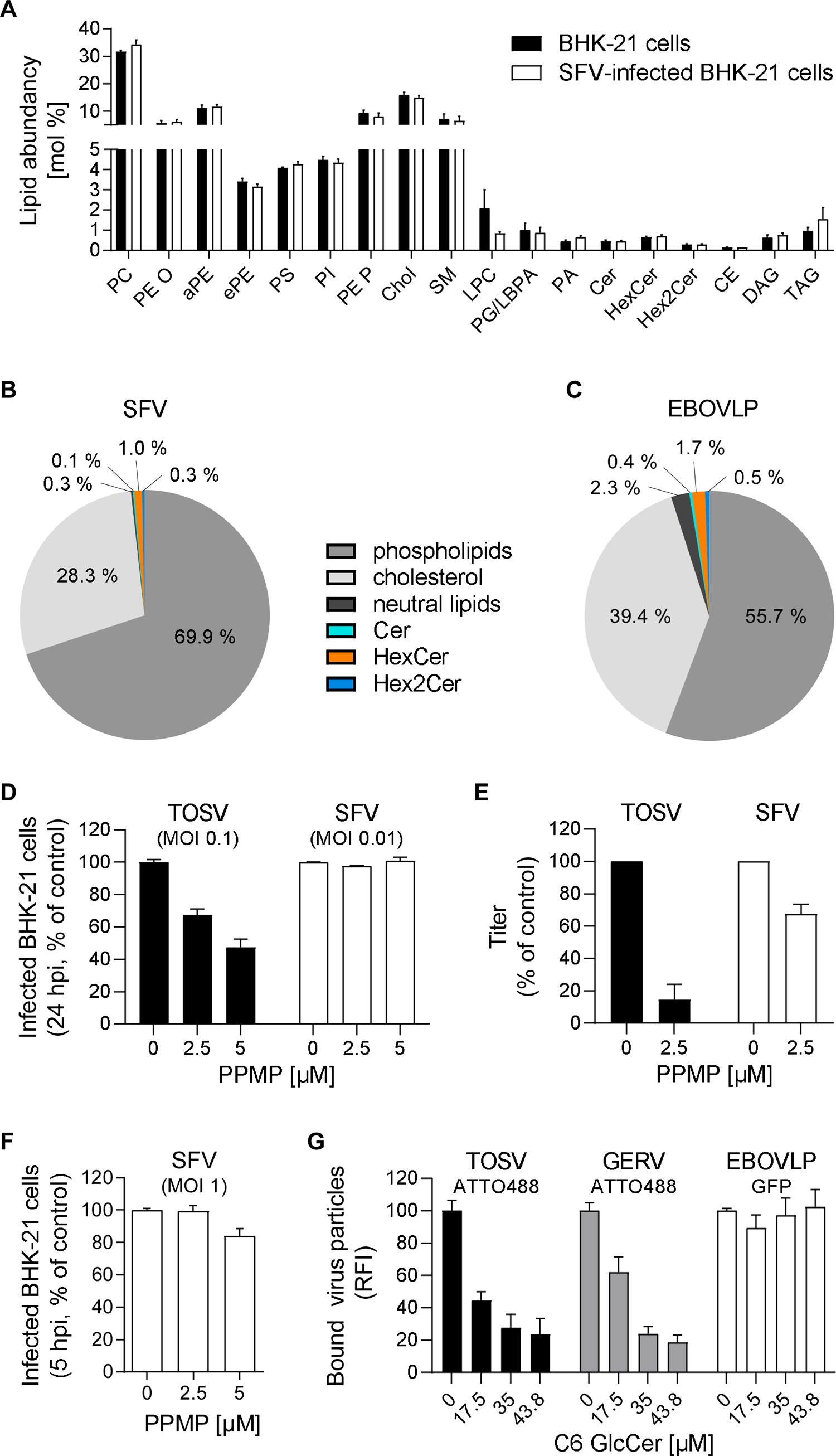
Glucosylceramide (GlcCer) is important for the attachment of viruses that bud from the Golgi. (A) BHK-21 cells were infected with Semliki forest virus (SFV) at a multiplicity of infection (MOI) of 0.01 and subjected to lipidomic analyses 14 h post-infection. (B) SFV particles were harvested 24 h post-infection and purified from the supernatant of infected cells and included in the analysis. (C) Ebola virus-like particles (EBOVLPs) were produced from HEK293T cells and purified prior to the lipidomic analysis with MS. (B-C) Phospholipids include all glycerophospholipids, sphingomyelin (SM), and lysophosphatidylcholine (LPC). The neutral lipids were cholesteryl ester (CE), diaculglycerol (DAG), and triacylglycerol (TAG). (D) BHK-21 cells were first treated with DL-*threo*-phenyl-2-palmitoylamino-3-morpholino-1-propanol (PPMP) at the indicated concentrations for 16 h and then infected with either TOSV (MOI ∼0.1) or SFV (MOI ∼0.01) in the continuous presence of PPMP. Infection was measured by flow cytometry 24 h later. (E) Infectious progeny of TOSV and SFV produced by BHK-21 cells in the presence of PPMP for 24 h was evaluated by focus-forming assay. The data are presented as the percentage of the control samples where viruses were produced in the absence of an inhibitor. (F) BHK-21 cells were pretreated with PPMP at the indicated concentrations for 16 h and exposed to SFV (MOI ∼1) for 5 h. Infected cells were immunostained for the SFV glycoprotein E2 and analyzed by flow cytometry. (G) Cells were preincubated with the indicated amounts of soluble C6-GlcCer for 2 h on ice and then exposed to ATTO488-labeled TOSV (TOSV-ATTO488) and ATTO488-labeled Germiston virus (GERV-ATTO488), and EBOVLPs containing green fluorescent protein (EBOVLP-GFP) on ice for 1 h. TOSV-ATTO488 and GERV-ATTO488 were used to infect cells at MOIs of 4 and 15, respectively. Approximately 500 ng of EBOVLP-GFP total protein was used to bind to 1×10^5^ cells. Virus binding was measured by flow cytometry, and the data were normalized to those in control samples processed in the absence of soluble C6-GlcCer. RFI, relative fluorescence intensity.

To evaluate the functional importance of GlcCer to human pathogenic phenuivirus infection, we extended our investigation to include TOSV. Similar to other phenuiviruses and bunyaviruses, TOSV buds into the Golgi network. BHK-21 cells were infected with TOSV in the presence of PPMP. We observed that the TOSV infectivity decreased with increasing concentrations of PPMP (Figure 7D). In addition, the release of infectious progeny was severely inhibited in the presence of PPMP (Figure 7E). In contrast, GlcCer depletion by PPMP had exerted no adverse effect on SFV infection or the subsequent production of infectious viral particles (Figure 7D to 7F). Finally, we assessed the capacity of soluble C6-GlcCer to inhibit TOSV binding. For this assay, purified stocks of TOSV were labeled with the fluorescent dye ATTO488, and then, the fluorescent virus (TOSV-ATTO488) was allowed to bind BHK-21 cells in the presence of increasing soluble C6-GlcCer concentrations on ice for 1 h. Soluble C6-GlcCer abolished TOSV binding (Figure 7G). Similar results were obtained when Germiston virus (GERV) was tested in a manner identical to TOSV (Figure 7G). GERV belongs to the *Peribunyaviridae* family in the *Bunyavirales* order and is thus related to phenuiviruses. Similar to phenuiviruses, peribunyaviruses bud and assemble in the Golgi network. As expected, EBOVLP binding was not affected by soluble C6-GlcCer (Figure 7G).

Collectively, the results suggested that viruses incorporate the glycolipid GlcCer when leaving the cell from the Golgi network but not from the plasma membrane, consistent with the transformation of Cer to GlcCer in this organelle. Overall, our study demonstrated that the glycolipid GlcCer mediates virus attachment to host cells, presumably through specific interactions with cellular receptors.

## Discussion

The lipid composition of enveloped viruses plays a major role in the structural organization of virions, which in turn contribute to virus binding and membrane fusion for efficient release of viral genomes into host cells. In this study, we evaluated the impact of a phenuivirus, *i.e.*, UUKV, on the lipidome of infected cells and determined the lipid composition of the envelope of newly produced viral particles. With the exception of HexCer, UUKV did not alter the lipid composition of infected cells. The strong enrichment of Chol, SM, and PE P in the viral envelope supports a budding of viral particles from post ER/post *cis* Golgi compartments. This is consistent with the accumulation of the UUKV glycoproteins Gn and Gc more in *trans* in the Golgi network, from where virions bud (Overby et al., 2006; Uckeley et al., 2019).

Our results clearly show that UUKV incorporates the glycolipid GlcCer in its envelope to attach to host cells. We also found that GlcCer promotes virus binding and infection by TOSV. These data are in agreement with studies showing that the infectious entry of the phenuivirus DABV relies on UGCG (Drake et al., 2017), the enzyme responsible for the synthesis of GlcCer from Cer in the Golgi (Ichikawa and Hirabayashi, 1998). Similar observations have recently been reported for the unrelated severe acute respiratory syndrome coronavirus 2, SARS-CoV-2 (Vitner et al., 2021). Although the GlcCer content of DABV and other phenuiviruses as well as SARS-CoV-2 remained to be determined experimentally, it is likely that many viruses that assemble in the Golgi apparatus can use GlcCer to infect host cells.

One phenuivirus, UUKV, was used to further analyze the role of glycolipids in the infection process. GlcCer was found to be enriched in infected cells. The reason is probably that GlcCer fails to undergo efficient transformation to Hex2Cer within the GSL synthesis pathway upon infection. UUKV, like other phenuiviruses, buds into the Golgi network and may interfere with the vesicular trafficking between the ER and Golgi network necessary for the processing of Cer into GSL (Ichikawa and Hirabayashi, 1998; Sun et al., 2018). Consistently, the release of UUKV infectious progeny relies on Golgi-specific brefeldin A-resistance guanine nucleotide exchange factor 1 (GBF1) (Uckeley et al., 2019), a guanine nucleotide exchange factor (GEF) that regulates ADP-ribosylation factor (ARF) and coat protein I (COPI)-dependent Golgi - ER trafficking (García-Mata et al., 2003). Alternatively, one or more of the four UUKV structural proteins that accumulate in the Golgi compartments to form new viral particles may specifically bind to HexCer or directly or indirectly regulate host cellular proteins that govern the GSL synthesis pathway.

Our lipidomic MS approach showed that HexCer not only accumulates in the infected cells but is also incorporated into the envelope of UUKV viral particles. The structures of GlcCer and GalCer are very similar, and our lipidomic MS analysis did not allow discrimination between the two types of HexCer molecules. The conversion from Cer to GalCer is ensured by GalCer synthases and limited to only a few tissues (Basu et al., 1968; Stahl et al., 1994). In contrast, Cer is primarily converted to GlcCer by UGCG in most cell types (Burger et al., 1996). Consistent with this, our subsequent analysis confirmed that GlcCer and probably not GalCer plays a role in UUKV infection.

Our results indicate that GlcCer was also embedded in both TOSV and GERV particles. GERV belongs to the *Peribunyaviridae* family, which is closely related to the *Phenuiviridae* family (Windhaber et al., 2022), and similar like phenuiviruses, peribunyaviruses bud into Golgi compartments (Elliott, 2014; Elliott and Brennan, 2014; Léger and Lozach, 2015). In contrast, SFV and EBOVLPs assemble at the plasma membrane, and our analysis showed no significant level of GlcCer level in their respective viral envelope. The lipid composition of other viruses that bud on the cell surface, such as vesicular stomatitis virus (VSV) (Patzer et al., 1979), has been analyzed, and GlcCer was not found in the VSV envelope (Kalvodova et al., 2009). Moreover, an excessive level of GlcCer has been shown to be detrimental to the cell cycle of influenza A virus (IAV) (Sillence et al., 2002; Drews et al., 2019; Drews et al., 2020). Although more viruses need to be analyzed to support these findings, GlcCer is likely incorporated only into the envelope of viruses that assemble in the Golgi.

Our data clearly showed a biological function for GlcCer in the viral envelope. The glycolipid contributed to both viral binding and infection. Soluble C6-GlcCer prevented UUKV, TOSV, and GERV attachment, but did not affect EBOVLP binding, consistent with the absence of GlcCer in the EBOVLP envelope. To our knowledge, so far only PS has been described to be associated with the viral envelope to facilitate the binding of different viruses, namely, Chikungunya virus, dengue virus, vaccinia virus, EBOV, and HIV (Callahan et al., 2003; Meertens et al., 2012; Jemielity et al., 2013; Moller-Tank et al., 2013; Kirui et al., 2021). However, PS significantly differs from GlcCer in several respects. For example, in contrast to GlcCer, PS is a negatively charged lipid abundant and widely exposed on the cell membrane surface. Recently, it has been shown that Chol interacts with viral fusion proteins and thereby facilitates the underlying fusion of the viral envelope with host cell membranes (Lee et al., 2021). In contrast, the GlcCer pathway appears to be finely and temporally regulated during UUKV infection; however, the details of the mechanisms remain to be discovered.

Because a high level of GlcCer in the UUKV envelope promotes viral binding, this glycosphingolipid likely interacts with one or more receptors on the host cell surface. Specifically, receptors may recognize the lipid core of GlcCer or the GlcCer molecules in the viral envelope may form larger clusters with other lipids in the plasma membrane of target cells, bringing the viruses close to receptor complexes. An alternative scenario suggests that the glucose moiety on ceramide is involved in interactions between viruses and host cell receptors. Macrophage inducible C-type lectin (Mincle) recognizes glucose residues and GlcCer, among other glycolipids (Nagata et al., 2017). In addition, this lectin has recently been shown to bind the phenuivirus RVFV and La Crosse virus, a peribunyavirus similar to GERV (Monteiro et al., 2019; Schön et al., 2022). Further investigations will be needed, not only to determine whether Mincle is an attachment factor or receptor for UUKV and other viruses carrying GlcCer molecules in their envelope, but also to identify the receptors that can capture enveloped viruses through GlcCer.

Significant efforts have been made to identify virus receptors. However, although thousands of viruses have been sequenced, only a few receptors have been described to date. Most researchers have focused on interactions between host cell surface factors and viral envelope glycoproteins or the carbohydrates they carry (Boulant et al., 2015). Lipids in viruses are difficult to investigate and only a handful of viral lipidomes have thus far been reported. Thus, the role of viral envelope lipids in infectious entry remains largely enigmatic. Our lipidomic analysis with MS of UUKV expanded the known virus lipidomes. By acting as a major molecular compound in the viral envelope that promotes virus binding to cells, the glycolipid GlcCer plays a critical role in UUKV-receptor interactions and viral tropism. We propose that, with the ability to incorporate glycolipids into their envelope, viruses can modulate tropism and increase interactions with host cells via glycan residues in their envelope, not solely through their viral glycoproteins. The results presented here offer new perspectives based on this novel virus binding modality for future receptor identification.

## Materials and Methods

### Cells

All products used for cell culture were obtained from Thermo Fisher Scientific or Merck. BHK-21 cells were grown in Glasgow’s minimal essential medium (GMEM) supplemented with 10% tryptose phosphate broth (TPB) and 5% fetal bovine serum (FBS). BHK-21 cells that stably express the human C-type lectin DC-SIGN were obtained by transduction with a TRIPΔU3 lentiviral vector encoding DC-SIGN as previously reported (Lozach et al., 2005). Human A549 lung and HEK293T kidney cells were cultured in Dulbecco′s Modified Eagle′s Medium (DMEM) supplemented with 10% FBS. The medium for culturing A549 cells was complemented with 1x nonessential amino acids (NEAAs). All mammalian cell lines were grown in an atmosphere in air with 5% CO2 at 37°C.

### Viruses and plasmids

UUKV strain 23 was obtained from plasmids and amplified in BHK-21 cells (Mazelier et al. 2016). The prototype strains of SFV, TOSV ISS, and GERV have been described previously (Pettersson and Kääriäinen, 1973; Helenius et al., 1980; Giorgi et al., 1991; Overby et al., 2006; Windhaber et al., 2022). All viruses were produced in BHK-21 cells and purified and titrated according to standard procedures (Meier et al., 2014; Léger et al., 2016; Hoffmann et al., 2018; Windhaber et al., 2022). Fluorescent labeling was performed with one molecule of viral glycoproteins per three molecules of ATTO488 dye (Atto-Tec) in 20 mM 4-(2-hydroxyethyl)-1-piperazineethanesulfonic acid (HEPES) following a protocol established by our group (Hoffmann et al., 2018). The MOI is presented on the basis of titers in BHK-21 cells. EBOVLPs were produced by transfecting HEK293T cells with plasmids encoding the EBOV structural proteins GP, VP40, and VP40-GFP at a 10:10:1 ratio. EBOVLPs were purified and concentrated as previously reported (Winter and Chlanda, 2021). Briefly, supernatants from transfected cells were first cleared through successive centrifugation cycles at 4°C. EBOVLPs were then pelleted in a 30%-sucrose cushion and resuspended in HNE buffer (10 mM HEPES, 100 mM NaCl, and 1 mM EDTA, pH 7.4). Residual sucrose was ultimately removed through a final ultracentrifugation step. The EBOVLP input was normalized based on the concentration of EBOV structural proteins as determined with a bicinchoninic acid (BCA) protein assay (Pierce™ BCA protein assay kit, Thermo Fisher Scientific).

### Small interfering RNAs (siRNAs), antibodies, and reagents

Nonoverlapping siRNAs against UGCG (siRNA UGCG 1: GUAAGAAACUGCUUGGGAA, UUCCCAAGCAGUUUCUUAC; siRNA UGCG 2: GGUUACACCUCAACAAGAA, UUCUUGUUGAGGUGUAACC) and siRNA negative control (scrambled: AGUAUUGAAUUUGCGACAA, UUGUCGCAAAUUCAAUACU) were designed by and obtained from Sigma–Aldrich. The mouse monoclonal antibody (mAb) 8B11A3 and the rabbit polyclonal antibodies (pAb) K1224 and K5 were directed against the UUKV nucleoprotein N and the glycoproteins Gn and Gc, respectively (Persson and Pettersson, 1991; Veijola and Pettersson, 1999). All these antibodies were kind gifts from the Ludwig Institute for Cancer Research, Stockholm, Sweden. The rabbit pAb U2, which has been described previously, recognizes the UUKV proteins N, Gn, and Gc (Lozach et al., 2011). The mouse mAb E2-1 targets the SFV glycoprotein E2 and was a generous gift from M.C. Kielian (Albert Einstein College of Medicine) (Kielian et al., 1990). Mouse immune ascitic fluid recognizes all TOSV structural proteins (and was a kind gift from R.B. Tesh, University of Texas) (Léger et al., 2016). The rabbit pAbs anti-actin C11, anti-UDP-UGCG, and anti-GlcCer were purchased from Sigma Aldrich, LSBio, and Antibody Research, respectively. The anti-DC-SIGN phycoerythrin-conjugated fragment antigen binding (Fab) 1621P was purchased from R&D Systems. Soluble C6-GlcCer was purchased from Cayman Chemical and dissolved in methanol. PPMP, PDMP, NB-DNJ, and NB-DGJ (Cayman Chemical) were also all dissolved in methanol. PPMP was assessed for cytotoxicity at the indicated concentrations with a CytoTox96 non-radioactive cytotoxicity colorimetric assay kit (Promega) according to the manufacturer’s instructions. Endo H and PNGase F were purchased from New England Biolabs and used according to the manufacturer’s recommendations.

### siRNA-mediated knockdown of UGCG

siRNA reverse transfections was performed with Lipofectamine RNAiMAX reagent according to the manufacturer’s protocol (Thermo Fisher Scientific) as reported previously (Uckeley et al., 2019). Briefly, 40,000 cells were transfected with siRNAs at final concentrations of 20 nM and seeded in a 24-well plate 3 days before infection.

### Flow cytometry-based infection assay

Flow cytometry-based assays were performed according to standard procedures (Lozach et al., 2011). Briefly, cells were infected with viruses at the indicated MOIs in the absence of serum for 1 h at 37°C. The viral input was replaced with serum-free medium, and the cells were incubated for up to 24 additional hours before fixation and permeabilization with 0.1% saponin. Infected cells were immunostained for detection of newly synthetized viral proteins with antibody 8B11A3 (anti-UUKV N), antibody 2E-1 (anti-SFV E2), and mouse ascitic fluid (TOSV) and quantified with a FACSCelesta cytometer (Becton Dickinson) and FlowJo software (TreeStar). For a UGCG inactivation assay, cells were pretreated with drugs at the indicated concentrations for up to 24 h at 37°C and then exposed to virus in the continuous presence of inhibitors.

### Lipidomic analysis with MS

To analyze the lipid content of infected cells and viral particles, BHK-21 cells were exposed to viruses in the absence of serum for the indicated incubation periods before fixation with 100% methanol. Viral particles were first purified by centrifugation with a 25%-sucrose cushion and then a 15-60% sucrose gradient. Sucrose was removed from the viral stocks by additional ultracentrifugation and washing steps in HNE buffer before fixation with methanol. For inactivation of UGCG with PPMP, cells were pretreated with 2.5 µM drug for 16 h before exposure to UUKV. Samples were subsequently subjected to lipid extraction in the presence of internal lipid standards and MS as previously described (Malek et al., 2021). Lipid extraction was performed using an acidic liquid–liquid extraction method (Bligh and Dyer, 1959), except for plasmalogens, which were extracted under neutral conditions. To ensure that similar amounts of lipids were extracted, a test extraction was performed to determine the concentration of PC as a bulk membrane lipid. Quantification was achieved by adding 1-3 internal lipid standards with a structure similar to that of the endogenous lipid species representing each lipid class. Sample volumes were adjusted to ensure that the lipid standard-to-lipid species ratios were in a linear range for quantification with the standard-to-species ratios within a range of >0.1 to <10. Following this approach, the relative quantification of lipid species was performed. Typically, 1,500-3,000 pmol of total lipid was used for lipid extraction. Lipid standards were added prior to extraction; the master mix consisted of 50 pmol PC (13:0/13:0, 14:0/14:0, 20:0/20:0, and 21:0/21:0, Avanti Polar Lipids), 50 pmol SM (d18:1 with semisynthesized N-acylated 13:0, 17:0, and 25:0 (Özbalci et al., 2013)), 100 pmol deuterated Chol (D7-Chol, Cambridge Isotope Laboratory), 30 pmol PI (17:0/20:4, Avanti Polar Lipids), 25 pmol PE and 25 pmol PS (both with semisynthesized 14:1/14:1, 20:1/20:1, and 22:1/22:1 (Özbalci et al., 2013)), 25 pmol DAG (17:0/17:0, Larodan), 25 pmol CE (9:0, 19:0, and 24:1, Sigma), and 24 pmol TAG (LM-6000/D5-17:0,17:1, and 17:1, Avanti Polar Lipids), 5 pmol Cer (d18:1 with semisynthesized N-acylated 14:0, 17:0, and 25:0 (Özbalci et al., 2013) or Cer (d18:1/18:0-D3, Matreya) and 5 pmol HexCer (d18:1 with semisynthesized N-acylated 14:0, 19:0, and 27:0, or GlcCer (d18:1/17:0, Avanti Polar Lipids)), 5 pmol Hex2Cer (d18:1 with an N-acylated C17 fatty acid chain), 10 pmol PA (17:0/20:4, Avanti Polar Lipids), 10 pmol PG (with semisynthesized 14:1/14:1, 20:1/20:1, and 22:1/22:1 (Özbalci et al., 2013)) and 5 pmol LPC (17:1, Avanti Polar Lipids). The PE P standard mix consisted of 16.5 pmol PE P-Mix 1 (16:0p/15:0, 16:0p/19:0, and 16:0p/25:0), 23.25 pmol PE P-Mix 2 (18:0p/15:0, 18:0p/19:0, and 18:0p/25:0), and 32.25 pmol PE P-Mix 3 (18:1p/15:0, 18:1p/19:0, and 18:1p/25:0). PE P was semisynthsized as previously described in (Paltauf and Hermetter, 1994). The final CHCl3 phase was evaporated under a gentle stream of nitrogen at 37°C. Samples were either directly subjected to MS analysis, or were stored at −20°C prior to analysis, which was typically completed within 1-2 days of extraction. Lipid extracts were resuspended in 10 mM ammonium acetate in 60 µl of methanol. Two-microliter aliquots of resuspended lipids were diluted 1:10 in 10 mM ammonium acetate in methanol in 96-well plates (Eppendorf twin tec 96) prior to measurement. For Chol measurements, the remaining lipid extract was evaporated again and subjected to acetylation as previously described in (Liebisch et al., 2006). Samples were analyzed on a QTRAP 6500+ mass spectrometer (Sciex) with chip-based (HD-D ESI Chip, Advion Biosciences) electrospray infusion and ionization Triversa Nanomate system (Advion Biosciences). In viral samples, TAG and DAG species were measured through a shotgun approach using a QExactive Plus instrument, and LipidXplorer software was used for data analysis as previously described in (Vvedenskaya et al., 2021). The MS settings and scan procedures are listed in Table S2. The data were evaluated by LipidView (Sciex) and an in-house-developed software (ShinyLipids). Sphingolipid species annotation is presented as <number of total C atoms in the sphingosine backbone and N-acylated fatty acid>:<number of hydroxyl groups>;<number of double bonds>, and for all other lipids, the <number of total C atoms in acyl or alkenyl/alkanyl>:<number of double bonds> is reported.

### Protein analysis

Cells were lysed with 0.1% Triton X-100, and lysates were diluted in lithium dodecyl sulfate (LDS) sample buffer (Thermo Fisher Scientific) and analyzed by SDS–PAGE (Nu-PAGE Novex 4-12% Bis-Tris gels; Thermo Fisher Scientific). Proteins were subsequently transferred to polyvinylidene difluoride membranes (iBlot Transfer Stacks; Thermo Fisher Scientific). The membranes were first blocked with intercept blocking buffer (LI-COR) and then incubated with primary antibodies against UGCG (LS-C107639, LS Bio), UUKV structural proteins, and actin (A2066, Sigma), all diluted in Tris-buffered saline containing 0.1% Tween (TBS-T) and intercept blocking buffer. After extensive washing, the membranes were incubated with the corresponding secondary antibodies conjugated to IRDye 680RD or 800CW (LI-COR). Proteins were analyzed with a LI-COR Odyssey CLx scanner and ImageJ v1.52p (NIH) software.

### Dot blot analysis

Cells were lysed in 0.1% Triton X-100, and lysates were added to nitrocellulose membranes using a Minifold^®^-1 Dot-Blot System (Whatman). The membranes were then immunostained with an antibody against GlcCer (111586, Antibody Research) following the procedure described in the Protein analysis section.

### Virus-binding assay

Viruses were allowed to bind to cells in binding buffer (DMEM or GMEM, pH 7.4, containing 0.2% BSA and 20 mM HEPES) on ice for up to 2 h, and binding was quantified by western blot or flow cytometry analysis after extensive washing. For analysis by western blotting, the amount of input virus was normalized to the amount of viral nucleoprotein N, which correlated with the number of particles, and bound viruses were detected with the rabbit pAb U2. For analysis by flow cytometry, at concentrations as high as 43.8 µM soluble C6-GlcCer was prebound to cells on ice for 1.5 h before fluorescent particles were added at the indicated MOIs, or GFP-containing EBOVLPs were added (500 ng per 10^5^ cells). Binding was analyzed with either a FACS Celesta, Canto, or Verse flow cytometer (Becton Dickinson) and FlowJo software (TreeStar).

### DC-SIGN identification at the cell surface

The location of DC-SIGN was assessed at the surface of BHK-21 cells (not permeabilized) by flow cytometry using an anti-DC-SIGN phycoerythrin-conjugated antibody (FAB1621P R&D Systems) according to a standard procedure (Lozach et al., 2005).

### Plunge freezing and cryo-electron tomography

To analyze UUKV particles produced in the presence or absence of PPMP, BHK-21 cells were pretreated either with 2.5 µM PPMP dissolved in methanol or with methanol for 16 h before exposure to UUKV at a low MOI (∼0.1). The supernatant was harvested 24 h post-infection, and the virus particles were purified first through a 25%-sucrose cushion and then through a 15-60% sucrose gradient. The remaining sucrose was removed by additional ultracentrifugation and washing steps with HNE buffer. UUKV virions were then inactivated with 4% paraformaldehyde for biosafety reasons prior to plunge freezing with a Leica GP2 plunger at 80% humidity and a 25°C air temperature. Viral suspensions were mixed with protein-A gold beads (10 nm), pipetted onto a 200 mesh copper grid coated with R2/1 Quantifoil carbon film, blotted from the backside of the mesh for 3 seconds with filter paper (Whatman No. 1), and plunge frozen in liquid ethane cooled to −183°C. A tilt series was with a Krios cryo-transmission electron microscope at 300 keV (Thermo Fisher Scientific) equipped with a K3 direct electron detector and Quanta Imaging Filter (Gatan) with an energy slit set to 20 eV. A dose-symmetric tilt series obtained at increments of 3° and a tilt range of 120° was acquired with SerialEM 4.0 (Mastronarde, 2005) at a magnification of 42,000 × (pixel size of 2.156 Å), defocus of −4 µm and electron dose per record of 3 e^-^/Å^2^. Projection images were aligned using fiducial gold and tomograms were reconstructed by weighted back projection in Etomo in the IMOD software package (Kremer et al., 1996) using SIRT-like filter 5 and a dose-weighting filter.

### Statistical analysis

Prism v9.1.1 (GraphPad Software) was used for plotting of numerical values in graphs and statistical analyses. The data are presented as the mean of at least three independent experiments ± standard error of the mean (SEM), unless stated otherwise in the figure legends. Figure legends indicate the statistical methods and p values when appropriate.

## Acknowledgments

This work was supported by CellNetworks Research Group funds and Deutsche Forschungsgemeinschaft (DFG) funding (grant numbers LO-2338/1-1 and LO-2338/3-1), INRAE starter funds, FINOVI AO14 (Fondation pour l’Université de Lyon), and the Agence Nationale de la Recherche (ANR) funding (grant number ANR-21-CE11-0012), all awarded to P.-Y.L. The authors also gratefully acknowledge the data storage service SDS@hd supported by the Ministry of Science, Research, and the Arts Baden-Württemberg (MWK) and DFG through grant INST 35/1314-1 FUGG and INST 35/1503-1 FUGG. We acknowledge Vibor Laketa and the Infectious Diseases Imaging Platform (IDIP) at the Center for Integrative Infectious Disease Research (CIID) Heidelberg, as well as the Cryo-EM Network at Heidelberg University (HD-cryoNet) for infrastructure access and support. PC and SW acknowledge Chica and Heinz Schaller foundation.

---

**Supplemental Figure S1.**
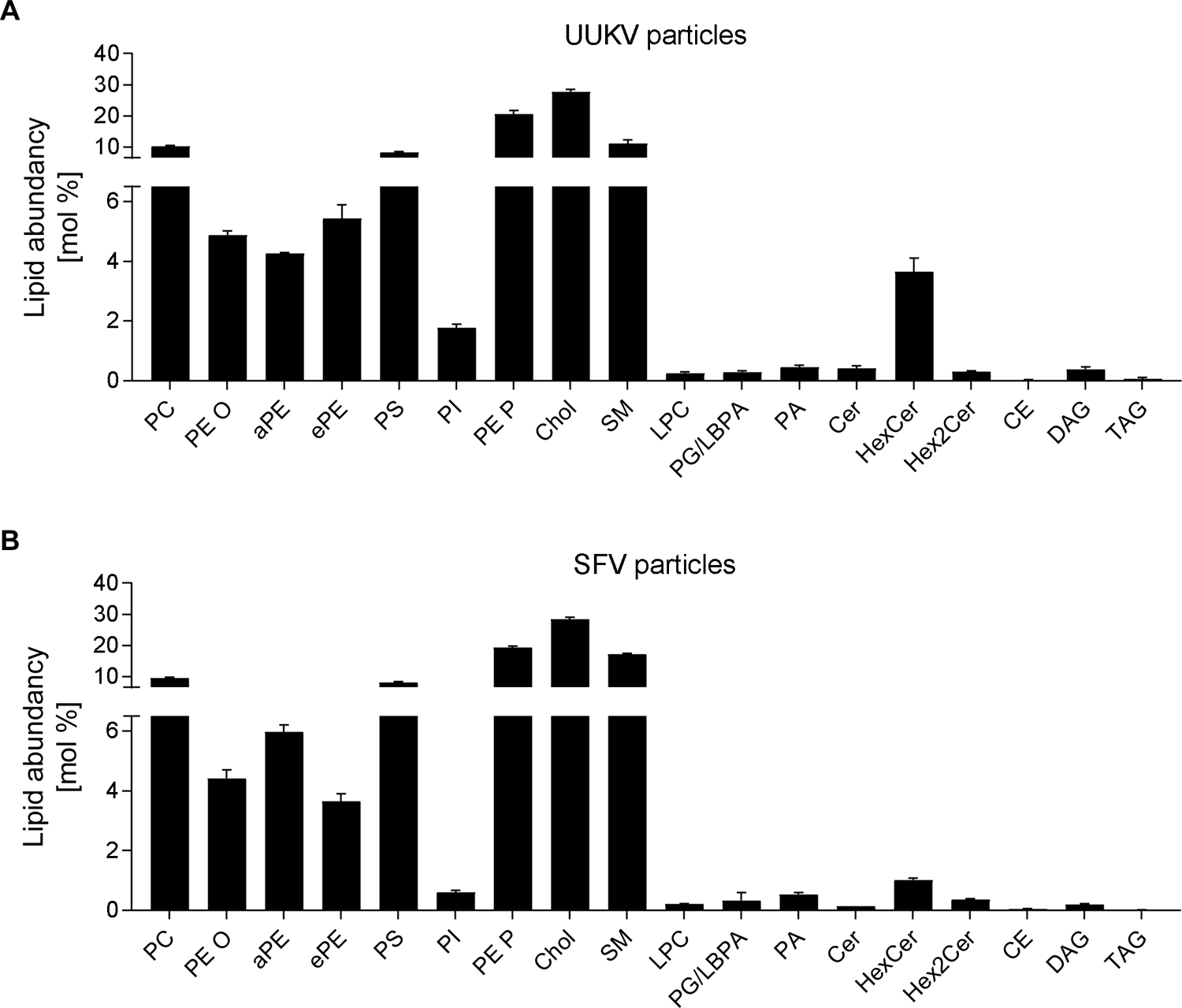
Mass spectrometry (MS) analyses of viral particles. Supernatant from infected BHK-21 cells was harvested 48 h post-infection (Uukuniemi virus (UUKV)) or 24 h post-infection (Semliki forest virus (SFV)), and UUKV particles (A) and SFV particles (B) were purified prior to quantitative MS-based lipid analysis.

**Supplemental Figure S2.**
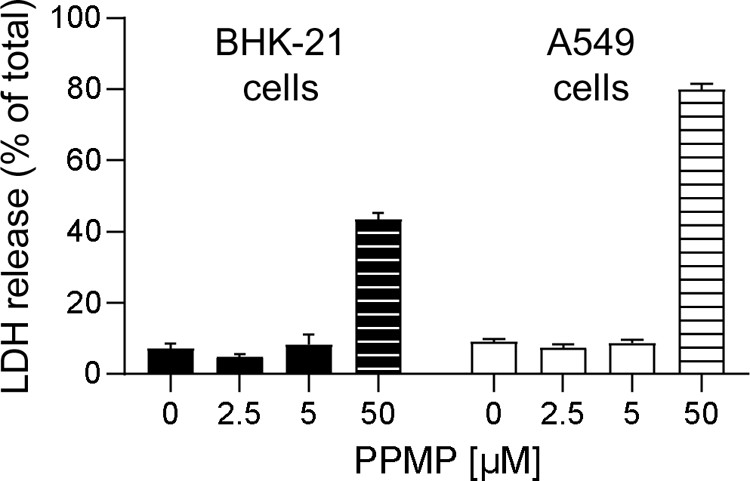
DL-*threo*-phenyl-2-palmitoylamino-3-morpholino-1-propanol (PPMP) cytotoxicity in A549 and BHK-21 cells. Assessment of the cytotoxicity of PPMP in the range of concentrations applied to A549 and BHK-21 cells, as determined with a CytoTox96 non-radioactive cytotoxicity colorimetric assay kit. Values were normalized to those of untreated cells after lysis, which corresponded to the maximum possible release of lactate dehydrogenase into the extracellular medium.

**Supplemental Figure S3.**
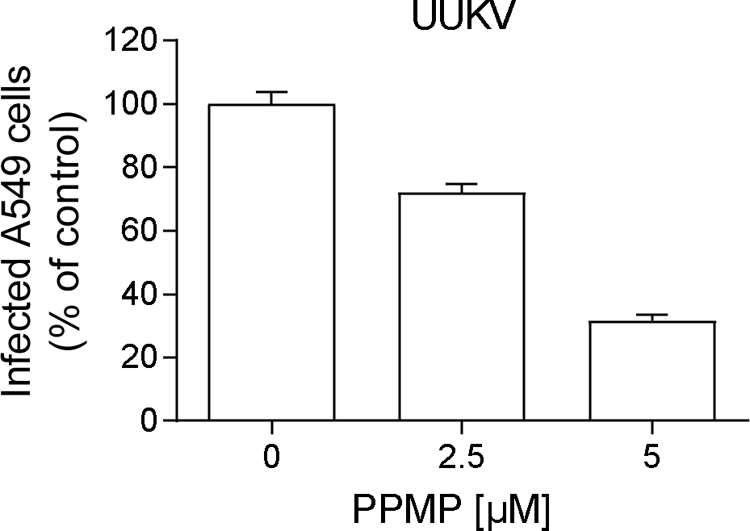
DL-*threo*-phenyl-2-palmitoylamino-3-morpholino-1-propanol (PPMP) treatment reduces UUKV infection in A549 cells. A549 lung epithelial cells were pretreated with PPMP for 16 h and then exposed to UUKV (multiplicity of infection ∼2) in the continuous presence of the inhibitor. Infected cells were harvested 8 h later and immunostained for UUKV nucleoprotein N. Infection was analyzed by flow cytometry, and the data were normalized to those of cells infected in the absence of the inhibitor; *i.e.*, it was reported as the percentage of the control.

**Supplemental Table S1.** Lipidomic analysis of BHK-21 cells infected with UUKV. This table is related to Figure 1A and 1D.

**Supplemental Table S2.** Lipidomic analysis of purified UUKV and SFV particles. This table is related to Figure 1C, 7B, S1A, and S1B.

**Supplemental Table S3.** Lipidomic analysis of UUKV-infected BHK-21 cells in the presence of DL-*threo*-phenyl-2-palmitoylamino-3-morpholino-1-propanol (PPMP). This table is related to Figure 3A and 3B.

**Supplemental Table S4.** Lipidomic analysis of BHK-21 cells infected with SFV. This table is related to Figure 7A.

